# Release of potential pro-inflammatory peptides from SARS-CoV-2 spike glycoproteins in neutrophil-extracellular traps

**DOI:** 10.1101/2020.05.02.072439

**Authors:** Aitor Blanco-Míguez, Borja Sánchez

**Affiliations:** Department CIBIO, University of Trento, Via Sommarive 9 38123, Povo, Trento, Italia; Department of Microbiology and Biochemistry of Dairy Products, Instituto de Productos Lácteos de Asturias (IPLA), Consejo Superior de Investigaciones Científicas (CSIC), Paseo Río Linares S/N 33300, Villaviciosa, Asturias, Spain; Ministry of Science, Innovation and University. Government of the Principality of Asturias. C/Trece Rosas nº 2 33005 Oviedo, Asturias, Spain

## Abstract

COVID-2019 has progressed in around 10-15% of patients to an acute respiratory distress syndrome characterized by extensive pulmonary inflammation and elevated production of pro-inflammatory cytokines. Neutrophil activation seems to be crucial in the initiation and perpetuation of this exacerbated lung inflammation. However, the precise mechanisms by which this activation occurs remain yet elusive. To this end, this *in silico* study tried to identify potential proinflammatory inducing peptides (PIPs) produced by the action of the elastase released in neutrophil-extracellular traps over SARS-CoV-2 particles. We found nine potential PIPs exclusive from the SARS-CoV-2, showing homology against T cell recognition epitopes. Moreover, 78 percent of these exclusive PIPs were found produced by the enzymatic cleavage on the spike glycoproteins, suggesting that high PIP concentrations might be released following SARS-CoV-2 huge replication rate. Therefore, these PIPs might play a role in the exacerbated inflammatory response observed in some patients.

**Highlights:**

- Nine potential PIPs were predicted exclusive from the SARS-CoV-2.
- SARS-CoV-2 PIPs showed homology against T cell recognition epitopes.
- Most of PIPs were produced by enzymatic cleavage of the spike glycoproteins.
- The release of these PIPs might be related to the increased inflammatory response observed in the patients.

**Graphical abstract:** 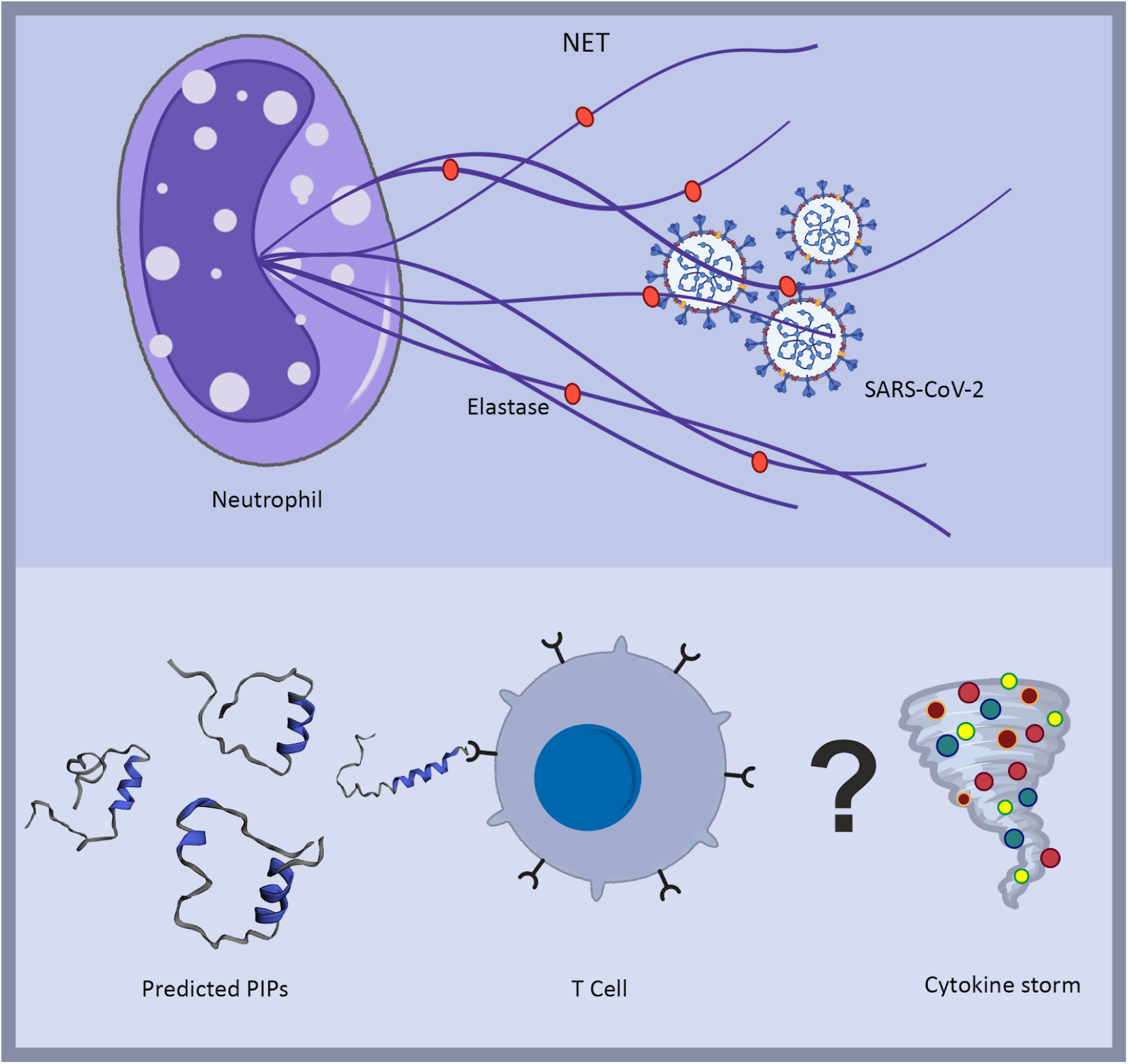

## Main text

Coronavirus disease 2019 (COVID-19) has progressed in around 10-15% of patients to an acute respiratory distress syndrome (ARDS) characterized by extensive pulmonary inflammation and elevated production of pro-inflammatory cytokines (Wang et al., 2020). Massive activation of pro-inflammatory macrophages and granulocytes -mostly neutrophils-, which involves secretion of high concentrations of IL1 and IL6, results in a cytokine storm that quickly evolves to ARDS (Shi et al., 2020). Activation of these immune cell sets seems to be key in ARDS development, although the precise mechanisms by which this activation occurs, including genetic susceptibility, remain yet elusive.

Recent efforts have been made in order to shed light over the potential factors inducing this aberrant immune response. Apart from macrophage/neutrophil activation, parallel analyses of CD4+ and CD8+ T cell subset activity strongly suggested that pro-inflammatory response may be intertwined with T cell activation that could prolong or even exacerbate ARDS (Ong et al., 2020). Potential B and T cell epitopes were also identified by sequence homology in the closely related SARS-CoV strains following an *a priori* epitope prediction, identifying promising targets that might develop strong immune responses against SARS-CoV-2 (Grifoni et al., 2020).

Neutrophil activation seems to be therefore crucial in the initiation and perpetuation of the exacerbated lung inflammation characterizing ARDS. Neutrophils possess a powerful strategy that serves as primary strategy against pathogens denominated neutrophil-extracellular traps (NETs). NETs are basically complex structures released from neutrophils containing genetic material and proteases that capture and kill microorganisms (Brinkmann et al., 2004), but also play an important role in sustaining inflammatory environments such as those observed in inflammatory bowel disease (Dinallo et al., 2019). An important NET component is elastase, a protease that acts over several proteins and substrates, mainly other proteases and extracellular matrix components. This protease is detectable at higher concentrations during pulmonary inflammation, and our hypothesis is that elastase can act over SARS-CoV-2 proteins releasing proinflammatory peptides.

To this end, a computational approach was proposed based on our knowledge predicting the immunomodulatory potential of microbial extracellular peptides (Blanco-Míguez et al., 2016). The proposed method combines an *in silico* digestion with machine learning and sequence alignment techniques (Figure 1). First, all SARS-CoV-2 proteins were *in silico* digested by the elastase action allowing one missed cleavage (see Methods). Then, we applied an ensemble learning strategy, combining 50 independent random forest models, including amino acid, dipeptide, composition–transition–distribution, physicochemical properties, and amino acid index, to predict the proinflammatory potential of the elastase-produced peptides (Manavalan et al., 2018) (Supplementary Table 1). To further check the capability of such peptides to induce a B / T Cell response, the 153 predicted proinflammatory inducing peptides (PIPs) were aligned against the potential B and T cell epitopes reported by Grifoni et al. (Grifoni et al., 2020) (see STAR methods). This led to a subset of sixteen PIPs, from which nine were found exclusive from the SARS-CoV-2 and five present also in the SARS-CoV, but not in other human coronaviruses, i.e. MERS-CoV, HCoV-229E, HCoV-NL63, HCoV-HKU1 and HCoV-OC43 (Table 1; Supplementary Table 2).

**Table 1.**
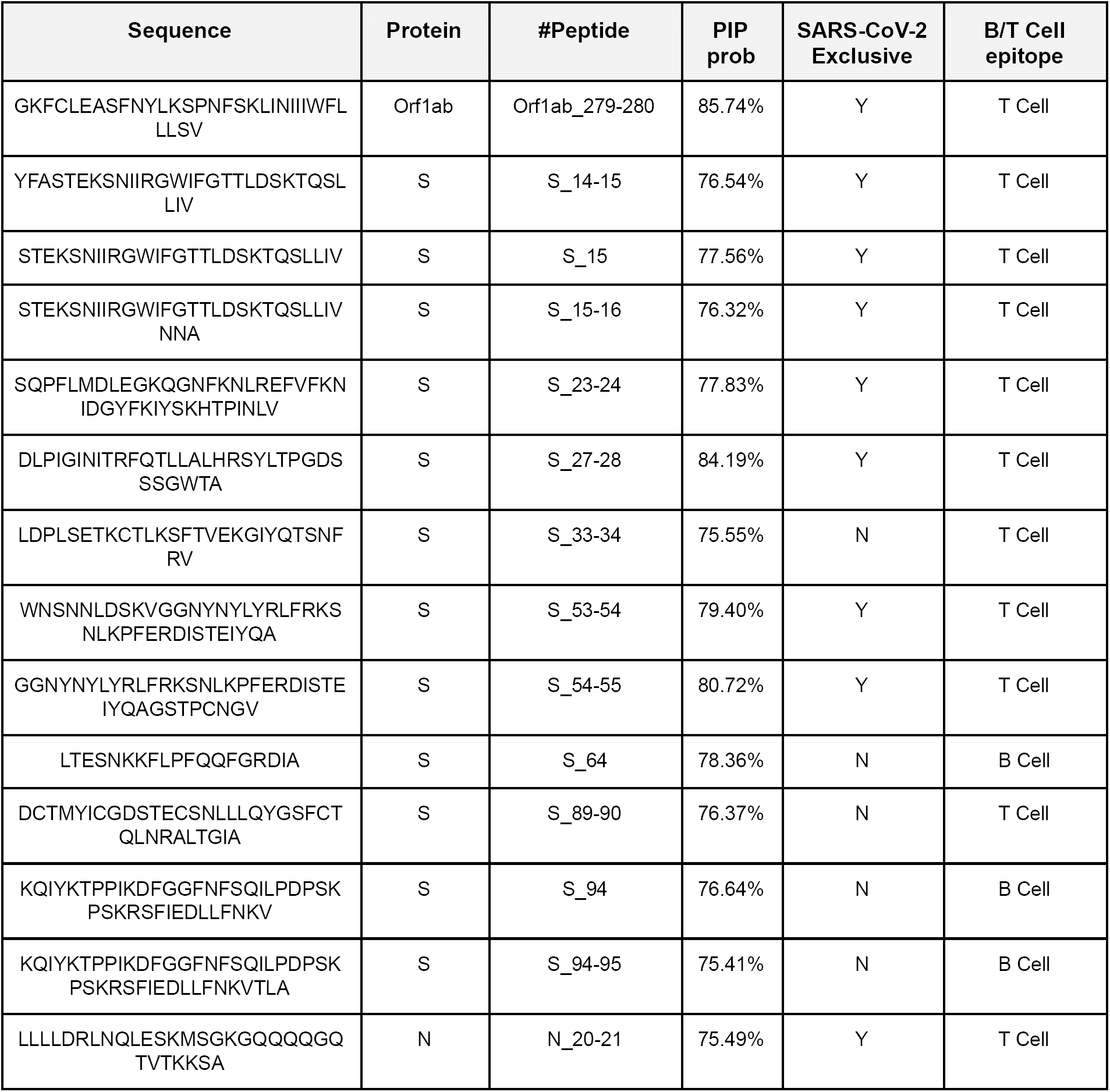
Predicted PIPs exclusive from the SARS coronavirus. S = spike glycoprotein; N = nucleocapsid phosphoprotein

**Figure 1.**
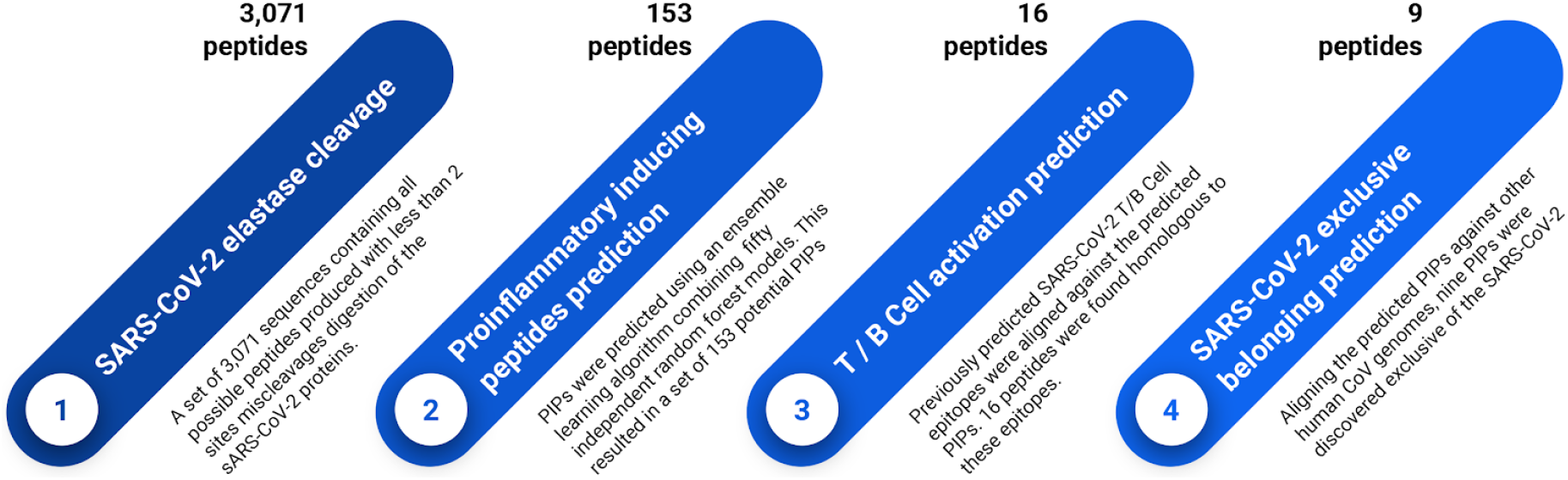
Computational pipeline to identify potential proinflammatory inducing peptides produced by the action of the elastase over the SARS-CoV-2 proteins.

Interestingly, all of the SARS-CoV-2 exclusive PIPs showed homology only against T-cell epitopes, agreeing with previous studies suggesting that the underlying mechanism behind the virus’ pro-inflammatory response was related to T cell activation (Ong et al., 2020). Moreover, 78 percent of these SARS-CoV-2 exclusive PIPs were found produced by the enzymatic cleavage on the spike glycoproteins, the second most abundantly expressed transcript of the virus (Kim et al., 2020). This finding suggests that high PIP concentrations might be released following SARS-CoV-2 huge replication rate from the outer and therefore accessible spike glycoproteins (Chu et al., 2020). High production of these spike PIPs might be related to the excessive pulmonary inflammation observed in the patients (Wang et al., 2020).

To our knowledge, this is the first study exposing the inflammatory potential of the peptides released after the action of the neutrophils in response to the SARS-CoV-2 infection. These results also suggest that the release of the PIPs could be related to increased inflammatory response on the human lungs. However, more investigation is needed to experimentally evaluate the action of these potential PIPs. We hope these findings would serve as a guide to design future investigations focused on generating more knowledge about ASDR triggers.

## Authors contributions

ABM and BS conceptualized the study and wrote the manuscript. ABM conducted the computational analyses.

## Declaration of interest

BS is on the scientific board and are co-founders of Microviable Therapeutics SL. The other authors have no competing interests.

## MATERIALS AND METHODS

The overall aim of the study was to identify potential proinflammatory inducing peptides (PIPs) produced by the action of the elastase over the SARS-CoV-2 proteins. To this end, all SARS-CoV-2 proteins were *in silico* digested by the elastase action allowing one miscleavage point as much. Then, an ensemble learning strategy was applied to predict the proinflammatory potential of the elastase-produced peptides. To further check the capability of such peptides to induce a B / T Cell response, the predicted PIPs were aligned against potential B and T cell epitopes. The resulting panel of PIPs were finally evaluated to check their exclusive presence in the SARS subtype.

### SARS-CoV-2 protein digestion

Protein digestion was performed over all the proteins of the reference SARS-CoV-2 isolate, Wuhan-Hu-1 (NCBI Reference Sequence: NC_045512.2). For this task, the RapidPeptidesGenerator version 1.1.0 (Maillet, 2020) was employed selecting the elastase as the digestion enzyme and disabling the miscleavages. Finally, in order to emulate all possible 1 site miscleavages situations, all the contiguous pairs of the previously digested peptides were joined together. This resulted in a set of 3,071 sequences containing all possible peptides produced with a perfect or a 1 site miscleavage digestion of the proteins.

### Proinflammatory inducing peptides prediction

The proinflammatory capability of the elastase-cleavaged peptides was assessed by applying an ensemble learning algorithm developed by Manavan et al (Manavalan et al., 2018). This algorithm combines 50 independent random forest models, including amino acid, dipeptide, composition–transition–distribution, physicochemical properties, and amino acid index, to predict potential proinflammatory inducing peptides. Unlike the original method, a PIP prediction threshold of 0.75 was selected to minimise the false positives.

### B and T cell response capability prediction

In order to check the capability of the predicted PIPs, they were aligned against the potential B and T cell epitopes reported by Grifoni et al (Grifoni et al., 2020) using BLASTp version 2.2.31 (Altschul et al., 1990). Parameters -word_size 1, -gapopen 9, -gapextend 1 and -evalue 200000 were selected. Successful alignments were defined by a 100% identity and a coverage greater than 80%.

### SARS exclusive belonging prediction

Exclusive belonging of the predicted PIPs to the SARS subtype was assessed aligning them against other the known human coronaviruses, i.e. MERS-CoV, HCoV-229E, HCoV-NL63, HCoV-HKU1 and HCoV-OC43 (NCBI Reference Sequences: NC_004718.3, NC_019843.3, NC_002645.1, NC_005831.2, NC_006577.2, NC_006213.1, respectively), using BLASTp version 2.2.31. Parameters -word_size 1, -gapopen 9, -gapextend 1 and -evalue 200000 were selected. Successful alignments were defined by a 60% identity and a coverage greater than 80%.

## Supplementary Material legends

Supplementary Table 1: Results of the ensemble learning prediction of proinflammatory inducing peptides over the elastase-cleavaged SARS-CoV-2 peptides.

Supplementary Table 2: Full report on the analyses of the predicted proinflammatory inducing peptides.

